# A conserved and druggable pocket in class B G protein coupled receptors for orally active small molecule agonists

**DOI:** 10.1101/2023.06.08.544171

**Authors:** Li-Hua Zhao, Qian He, Qingning Yuan, Yimin Gu, Xinheng He, Junrui Li, Kai Wang, Yang Li, Jianhua Shen, H. Eric Xu

## Abstract

Class B G protein-coupled receptors (GPCRs), including glucagon-like receptor 1 (GLP-1R) and parathyroid hormone receptor 1 (PTH1R), are peptide hormone receptors and important drug targets. Injectable peptide drugs targeting class B GPCRs have been developed for the treatment of many diseases, including type 2 diabetes, obesity, and osteoporosis, but orally available small molecule drugs are hotly pursued in the field, especially small molecule agonists of GLP-1R and PTH1R. Here we report the first high-resolution structure of the human PTH1R in complex with the stimulatory G protein (G_s_) and a small molecule agonist, PCO371, which reveals an unexpected binding mode of PCO371 at the interface of PTH1R and G_s_. The binding site of PCO371 is totally different from all binding sites previously reported for small molecules or peptide ligands in GPCRs. Residues that make up the PCO371 binding pocket are mostly conserved in class B GPCRs and a single mutation in PTH type 2 receptor (PTH2R) and two residue mutations in GLP-1R convert these receptors to respond to PCO371 activation. Functional assays reveal that PCO371 is a G-protein biased agonist that is defective in promoting PTH1R-mediated arrestin signaling. Together, these results uncover a distinct binding site for designing small molecule agonists for PTH1R and possible other members of class B GPCRs and define a receptor conformation that is only specific for G protein activation but not arrestin signaling. These insights should facilitate the design of distinct types of class B GPCR small molecule agonists for various therapeutic indications.

## Introduction

Class B G protein-coupled receptors (GPCRs) are peptide hormone receptors that are drug targets for many diseases, including osteoporosis, type 2 diabetes, obesity, bone metabolism diseases, cardiovascular disease, migraine, and depression^1–4^. Structures of all 15 members of class B GPCRs with peptide agonists have been determined in recent years^2^, providing important molecular mechanisms of hormone recognition and receptor activation for the whole class B GPCRs and rational templates for designing better peptidic and small-molecule drugs^2^. Class B GPCRs are different from class A GPCRs because many therapeutic small molecule agonist drugs have been developed for class A but not for class B GPCRs^5^. For class B GPCRs, despite great efforts toward discovering orally available non-peptidic agonists, few small molecule agonists of class B GPCRs are known^6, 7^. This is a difficult problem in class B GPCRs, because their natural ligands are peptide hormones, which activate the receptor through peptide binding to a large open pocket within the receptor transmembrane domain (TMD) and the high affinity binding of peptide hormones requires the interaction with the receptor extracellular domain (ECD)^8^. To date, only a few small molecule agonists of GLP-1R and PTH1R have been reported^9–17^. Several structures of GLP-1R with a partial or full non-peptidic small molecule agonist have also been reported^9, 13, 18–20^, which reveals that they bind to the same binding site of peptide hormones or to an allosteric site at the cytoplasmic end of TM6^9, 11, 13, 18, 21^. Nonetheless, there is no orally available small molecule drugs of class B GPCRs. It is challenging but remains a long-term goal to replace the injectable peptide drugs with oral drugs, with aims to improve the quality of life of patients, the profiles of side-effects, and the costs of peptide drugs.

Parathyroid hormone receptor 1 (PTH1R) is a classic member of class B GPCRs that regulates calcium homeostasis and skeleton development through activation by two endogenous peptide hormones, parathyroid hormone (PTH) and PTH-related peptide (PTHrP)^8, 22, 23^. PTH1R is a clinically proven target for hypoparathyroidism and osteoporosis, which can be treated with injections of PTH or PTHrP analogs ^8, 17^.

Recently Nishimura et al^24^ reported a human PTH1R agonist, PCO371, as a potent and orally available small molecule agonist that is currently being evaluated in a phase 1 clinical study for the treatment of hypoparathyroidism^17, 25^. However, the molecular mechanism of PTH1R activation by PCO371 remains unknown. In this paper, we report the structure of PTH1R bound to PCO371 and its functional characterization as a G-protein biased agonist. To our surprise, the structure reveals that PCO371 binds to an unexpected site at the interface between PTH1R and G-protein, distinct from all other sites known for GPCR ligands. Importantly, the PCO371 pocket is mostly conserved in class B GPCRs, thus opening a new avenue for designing small drug molecules targeting specifically to this pocket.

## Results

### Characterization of PCO371 and structure determination

PCO371 is a potent and orally available small molecule for treatment of hypoparathyroidism^24^. PCO371 was characterized as an agonist of PTH1R as it can induce cAMP production to the same level as did PTH (1–34)^17^, but it remains unknown whether PCO371 could induce PTH1R-mediated β-arrestin signaling (Fig. 1a). We first investigated their effects on G protein signaling pathways using cAMP accumulation, and their effects on β-arrestin signaling using β-arrestin recruitment assay. We confirmed that PCO371 is a full G-protein agonist, but discovered that PCO371, unlike PTH peptide, is defective in promoting PTH1R-mediated arrestin signaling (Fig. 1b-d). These data suggest that PCO371 is a G-protein biased agonist.

**Fig. 1.**
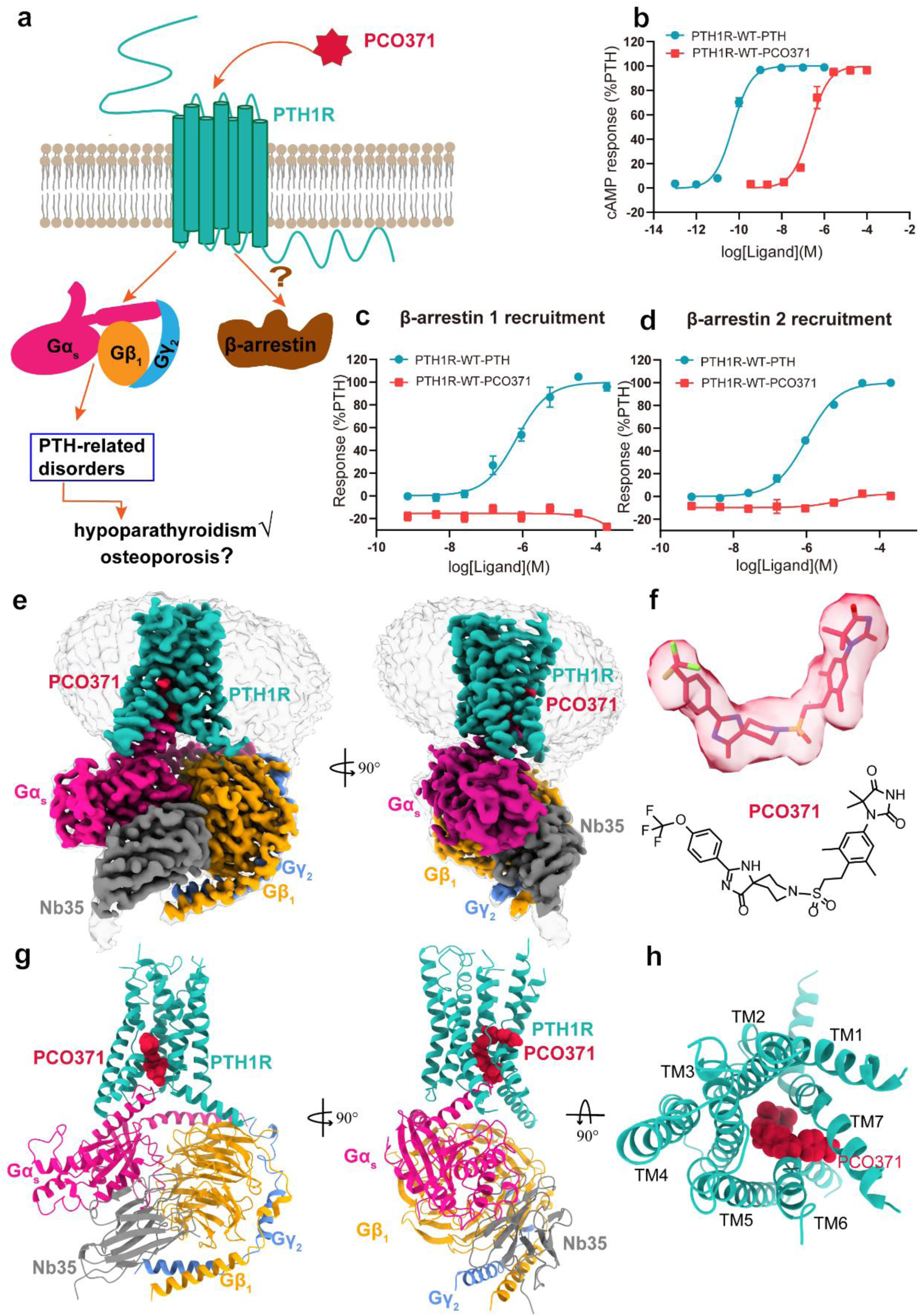
| Cryo-EM structure of G_s_-coupled PTH1R bound to PCO371. (**a**) PCO371 induced PTH1R signaling and potential pharmacological effects. **(b-d**) Concentration-dependent response curves of PCO371 to induce cAMP accumulation **(b**) and β-arrestin recruitment (**c-d**). Data were presented and graphed as means ± S.E.M. of three independent experiments, and each experiment was performed in triplicate. The data were normalized according to the maximal response of PTH. (**e**) Cryo-EM maps of PCO371-PTH1R-G_s_ complex. (**f**) Chemical structure of PCO371. (**g**) Cryo-EM structure model of PTH-PTH1R-G_s_ complex. (**h**) The top view shows the binding site of PCO371.

To study the G-protein-biased agonism of PCO371, we prepared the PCO371-bound PTH1R-G_s_ complex using the NanoBiT tethering strategy, which details are described in methods^26, 27^. The carboxyl terminus of PTH1R was truncated to residue H502 to increase the expression level of PTH1R as we showed previously (Extended Data Fig. 1a)^28^. The complex was purified by size-exclusion chromatography and verified by SDS gel (Extended Data Fig. 1b). The structure of PCO371-PTH1R-G_s_ complex was solved by cryo-EM to high resolution of 2.57 Å (Fig. 1e, Extended Data Fig. 2, and Extended Data Table 1). The high resolution cryo-EM map is sufficiently clear to place the receptor, the G_s_ heterotrimer, and the small molecule agonist in the PTH1R-G_s_ protein complex (Fig. 1e-h and Extended Data Fig. 3). Unlike the peptide-bound PTH1R-G_s_ structures, the PTH1R ECD was invisible in this PCO371-bound PTH1R-G_s_ structure due to the flexibility of the ECD in the absence of the peptide binding.

**Fig. 2.**
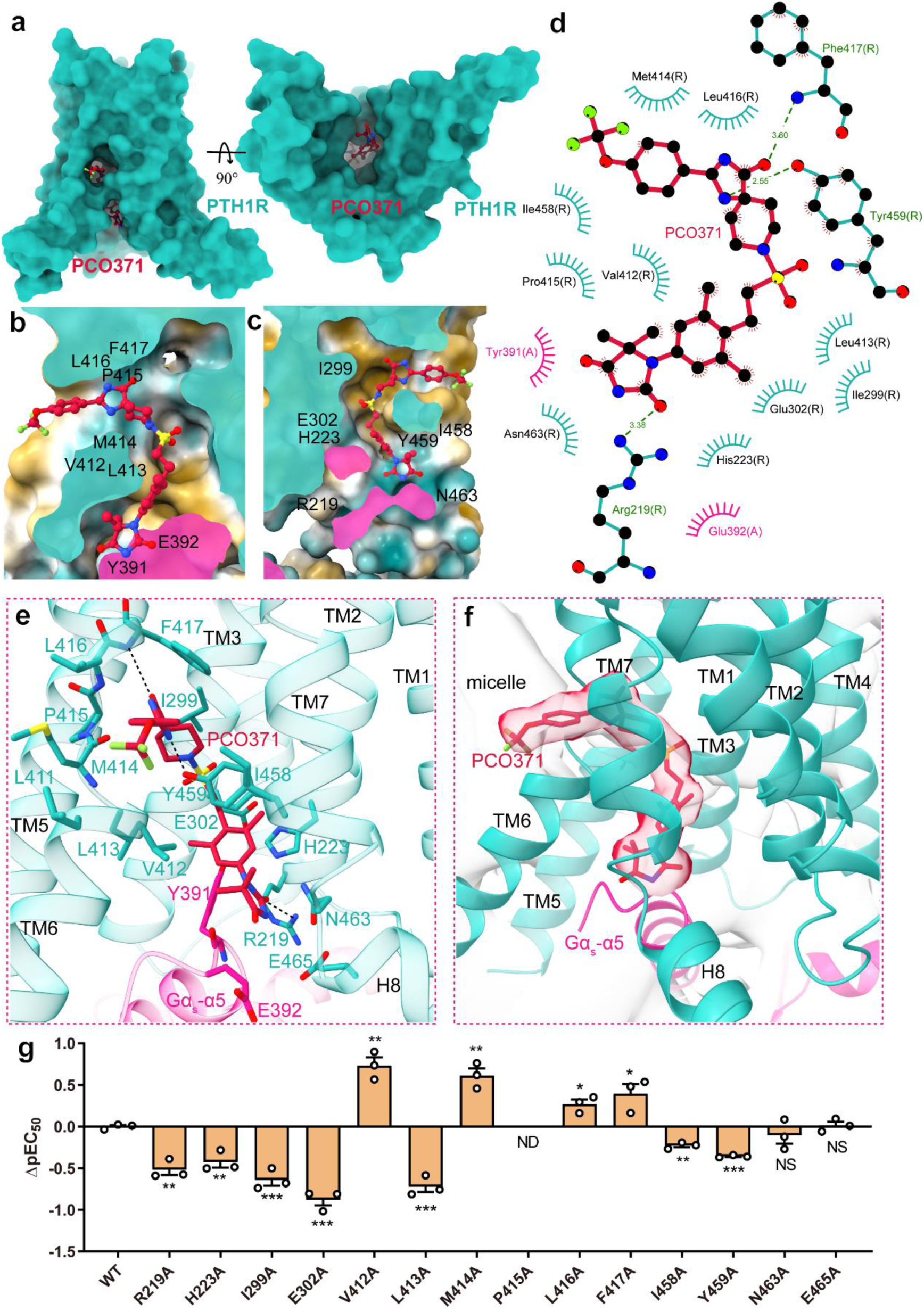
| Interactions of PCO371 with PTH1R. (**a**) The PCO371-binding pocket of PTH1R viewed from the side view and intracellular side. (**b-c**) Cross-section of the PCO371-binding pocket in PTH1R. (**d**) Interacting residues predicted by LigPlot using the full-length model. (**e**) Detailed interactions of PCO371 with residues in the binding pocket. (**f**) The bound PCO371 at the interface between PTH1R and G_s_ protein and the tail phenyl inserts into the detergent micelle. (**g**) Signaling profiles of PTH1R mutants of key residues on PCO371-induced cAMP accumulation. ΔpEC50 represents the difference between pEC50 values of the wild-type (WT) and the mutated PTH1Rs. Data from three independent experiments, each of which was performed in triplicate, are presented as mean ± SEM. Statistical differences between WT and mutations were determined by two-sided one-way ANOVA with Tukey’s test. \**P*<0.05; \*\**P*<0.01; \*\*\**P*<0.001 vs. WT receptor, ND, not detectable. NS, no significant difference. All data were analyzed by two-side, one-way ANOVA with Tukey’s test.

**Fig. 3.**
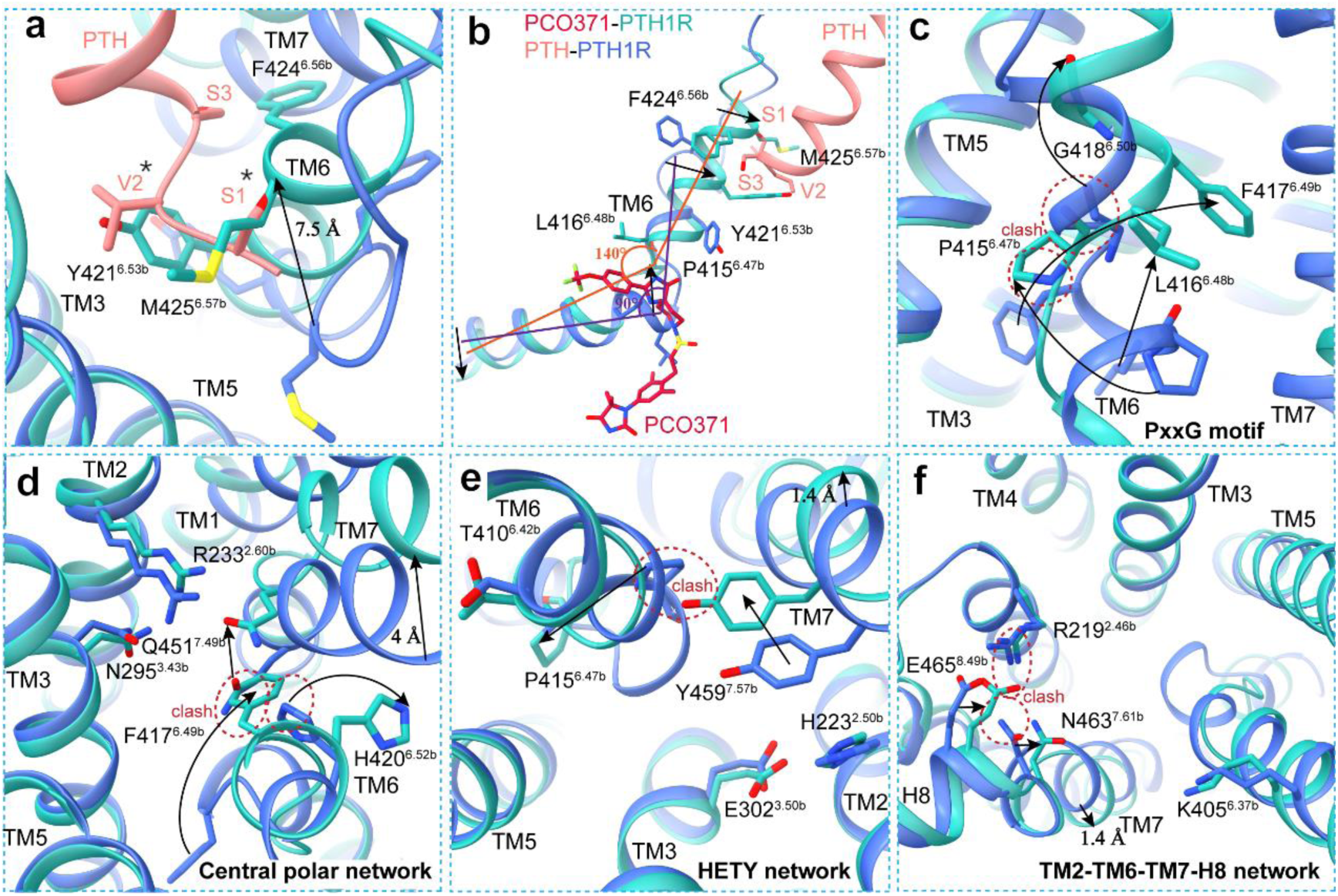
| Conformational changes of TMD helix bundles during receptor activation between PTH-bound and PCO371-bound PTH1R. (**a**) Structural comparison of the TMD bundles of the active PTH1R (light sea green) with PCO371 (crimson), PTH1R (royal blue) with PTH (light coral) (PDB: 8HA0). Hormone peptide, PCO371, G protein and Nb35 are omitted for clarity. (**b**) Comparison of TM6 conformational changes between the PCO371-bound and peptide-bound PTH1R structures. (**c-f**) Different conformations are shown for conserved residues and motifs in the active PTH1R, including the conserved PxxG motif (P415^6.47b–^L416^6.48b–^F417^6.49b–^G418^6.50b^), the central polar network (R233^2.60b–^N295^3.43b–^H420^6.52b–^Q451^7.49b^), the HETY network (H223^2.50b–^E302^3.50b–^T410^6.42b–^Y459^7.57b^) and the TM2-TM6-TM7-H8 network (R219^2.46b–^K405^6.37b–^N463^7.61b–^E465^8.49b^).

### Overall architecture

The overall structure of PTH1R exhibits a canonical seven-transmembrane domain fold of GPCRs and the hallmark of class B GPCRs activation with a kink in the middle of the TM6 (Extended Data Fig. 4a and b). We also observed several remarkably distinct features in PCO371-PTH1R-G_s_ structure compared to three cryo-EM structures of PTH1R in complexes with PTH, PTHrP, and LA-PTH as previously reported (Extended Data Fig. 4, Extended Data Fig. 5)^28, 29^. The notable difference is that PCO371 occupies a distinct ligand-binding pocket of PTH1R, comprising of intracellular portion of TM2, TM3, TM6 and TM7 as well as helix H8, at the interface between PTH1R and the G protein (Fig. 1h, Extended Data Fig. 4a and b). This binding pocket is different from the peptide hormone binding pockets of class B GPCRs and the small-molecule binding pockets of GLP-1R (Fig. 2, Extended Data Fig. 4b, 5, 6). In responding to PCO371 binding, the extracellular tips of helices TM1, TM6, and TM7 in the PCO371-PTH1R-G_s_ structure shift counterclockwise by as much as 7–8 Å, relative to their positions in the PTH-PTH1R-G_s_ structure (Extended Data Fig. 4c), which results in a collision between the extracellular end of TM6 and the bound PTH peptide, consistent with the report that the presence of PCO371 would inhibit the binding of PTH to its TMD^17^. On the other hand, relative to the peptide-bound PTH1R structures, we observed a ∼4 Å inward shift at the cytoplasmic end of TM6 as measured by the Cα of R400^6.32b^ and a 1.4 Å outward shift at the cytoplasmic end of TM7 as measured by the Cα of I458^7.56^ (Extended Data Fig. 4d). Together, these observations suggest that PCO371 induced a very distinct PTH1R conformation, unseen for structures of all other class B GPCRs, to couple with downstream signal transducers.

**Fig. 4.**
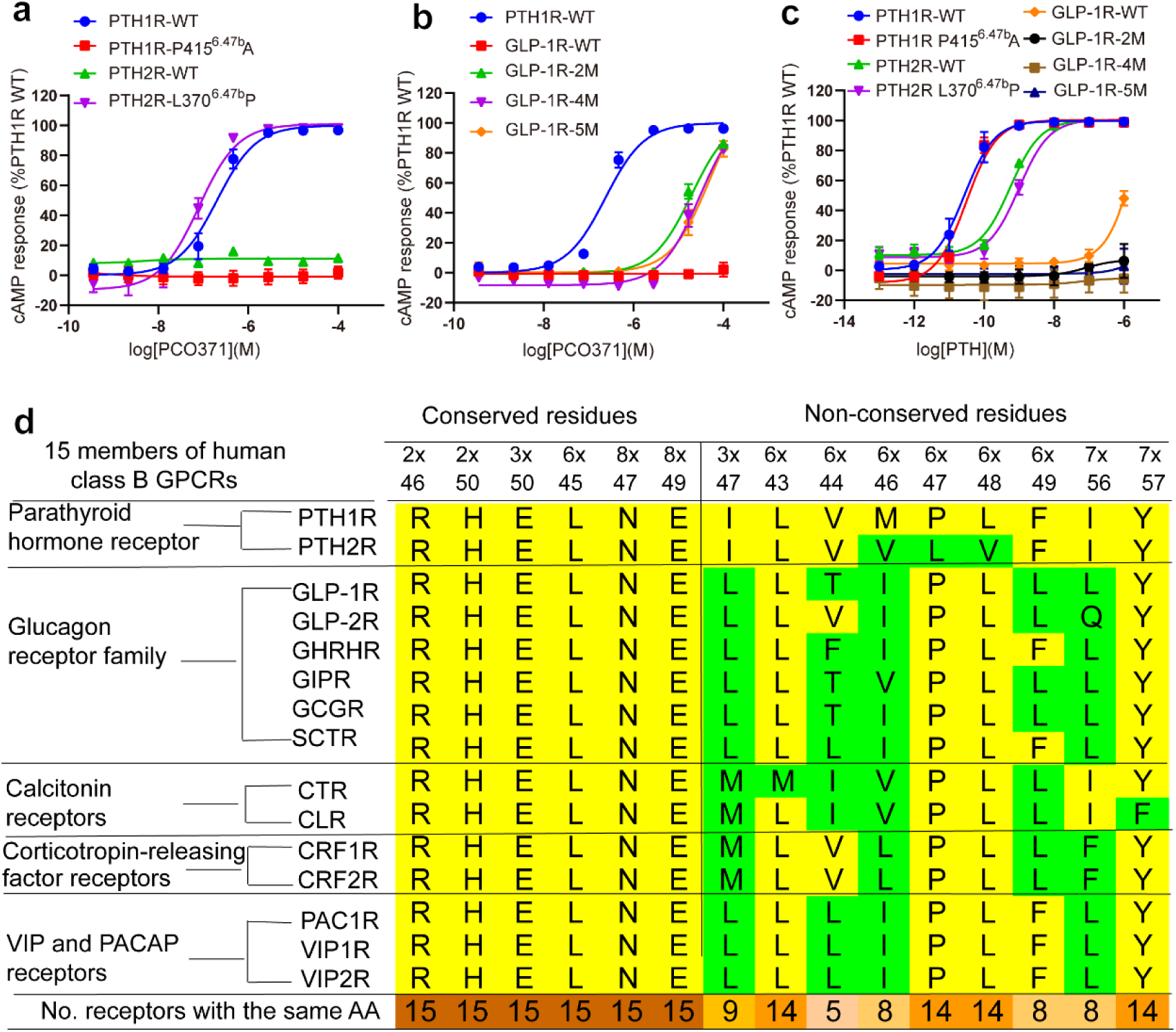
| Selectivity of PCO371 for PTH1R and the conservation of the PCO371-binding site in class B GPCRs. (**a** The cAMP production stimulated by PCO371 in the wild-types (WTs) and mutants of PTH receptors. (**b**) Stimulation of cAMP production by PCO371 in the WT and mutants of GLP-1R. Data from three independent experiments (n=3), each of which was performed in triplicate, are presented as mean ± SEM. (**c**) Stimulation of cAMP production of wildtype or mutated PTH1R, PTH2R and GLP-1R by PTH. Data from three independent experiments (n=3) performed in technical triplicate are presented as mean ± SEM. GLP-1R-2M, GLP-1R-4M and GLP-1R-5M are the combined mutations of two residues (L244^3.47b^I and L360^6.49b^F), four residues (L244^3.47b^I/T355^6.44b^V/L360^6.49b^F/L401^7.56b^I), and five residues (L244^3.47b^I/T355^6.44b^V/L360^6.49b^F/L401^7.56b^I/N407^8.48b^G). (**d**) Sequence alignment of conserved and non-conserved residues forming the pocket of PCO371 in class B GPCRs.

**Fig. 5.**
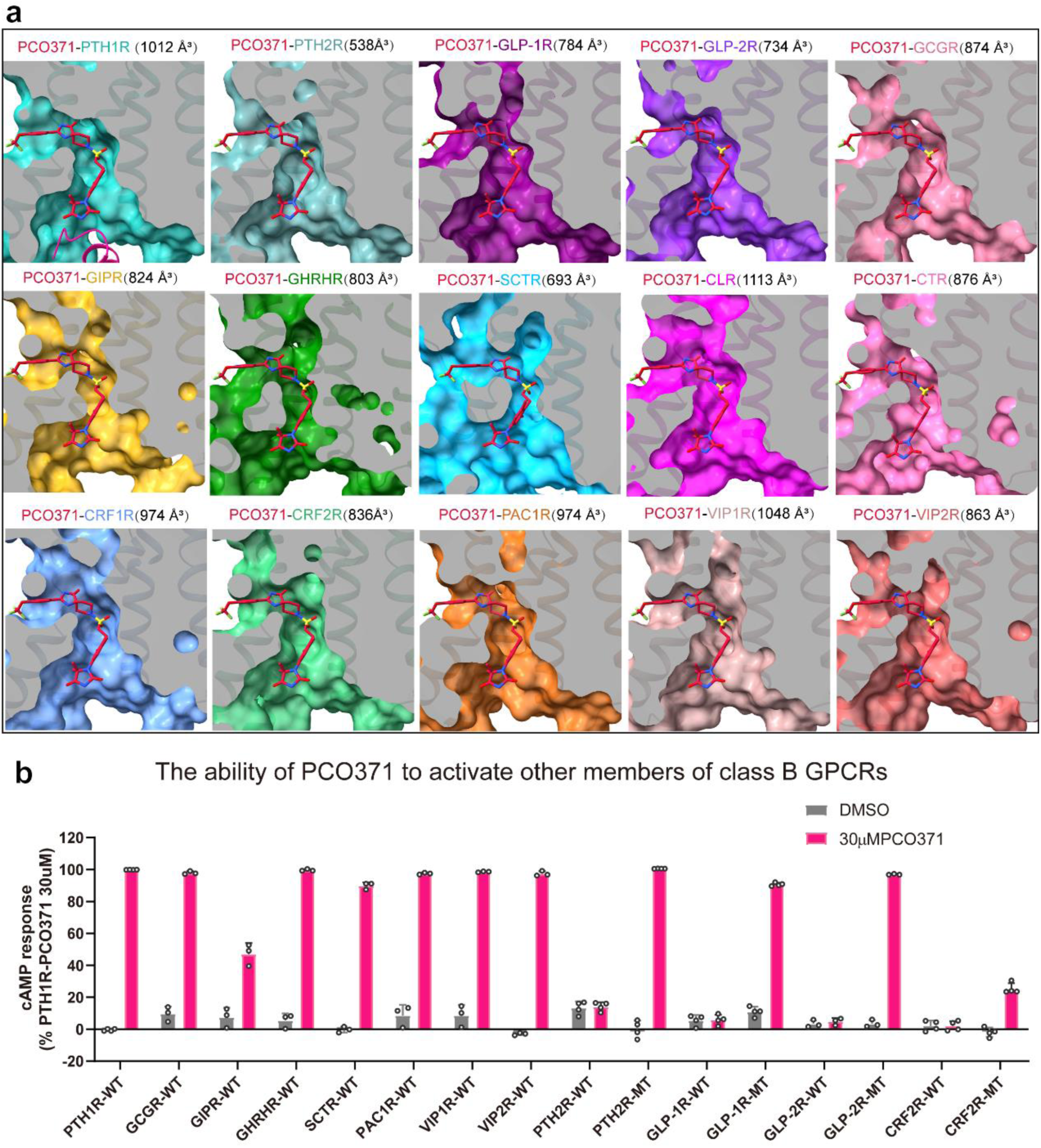
| A mostly conserved PCO371-like binding pocket in class B GPCRs. (**a**) The PCO371-like binding pocket is mostly conserved in other members of class B GPCRs by structural modeling. The volume calculation shows these pockets in different receptors are similar in all class B GPCR receptors. Peptides, G protein and Nb35 are omitted for clarity. PDB: 7F16, PTH2R: cadet blue; PDB: 6X1A, GLP-1R: purple; PDB: 7D68, GLP-2R: blue violet; PDB: 7CZ5, GHRHR: green; PDB: 7DTY, GIPR: goldenrod; PDB: 6WPW, GCGR: pale violet red; PDB: 6WZG, SCTR: deep sky blue; PDB: 6NIY, CTR: hot pink; PDB: 6E3Y, CLR(CGRPR): magenta; PDB: 6PB0, CRF1R: cornflower blue; PDB: 6PB1, CRF2R: medium sea green; PDB: 6P9Y, PAC1R: chocolate; PDB: 6VN7, VIP1R: rosy brown; PDB: 7VQX, VIP2R: Indian red. (**b**) PCO371 has pan-agonist activity in wildtype and mutated class B GPCRs. The mutated receptors have two corresponding mutations as GLP-1R that regain response to PCO371.

### PCO371 has an unanticipated binding pattern

Within the structure, PCO371 adopted a horizontal ‘‘U’’-shape pose that wraps around the bottom half (intracellular half) of TM6, (Fig. 2a-c), forming extensive interactions with residues within TM2, TM3, TM6, TM7 and H8 of receptor and the α5 helix of Gα_s_ (Fig. 2d). The chemical structure of PCO371 is comprised of the head imidazolidinone, the middle dimethylphenyl, the sulfonamide linker, the piperidine motif, the middle spiro-imidazolone, and the tail trifluoromethoxy phenyl (Fig. 1f and Extended Data Fig. 7a). The head imidazolidione and the middle phenyl of PCO371 are embedded in the interface between the receptor and the Gα_s_ protein and form interactions with the residues within TM2, TM6, TM7 and H8, as well as with α5 helix of Gα_s_ (Figure 2d-e). The head imidazolidione of PCO371 also forms a hydrogen bond with R219^2.46b^ and a polar interaction with Y391 from α5 helix of Gα_s_. In addition, the head imidazolidione and the middle phenyl of PCO371 form extensive hydrophobic interactions with the receptor and α5 helix of Gα_s_ (Fig. 2d-e). Specifically, both Y459^7.57b^ and Y391 of Gα_s_-α5 form pi stacking interactions with the middle phenyl of PCO371. (Fig. 2d-e).

In the middle of PCO371, the sulfonamide group forms polar interactions with E302^3.50b,^ the piperidine group form hydrophobic contacts with I299^3.47b^ (Fig. 2d-e). The middle spiro-imidazolone group of PCO371 forms hydrogen bond interactions with the main chain amine of F417^6.49b^ and side chain of Y459^7.57b^. The middle spiro-imidazolone together with the tail phenyl group form extensive hydrophobic interactions with PTH1R residues from TM3, TM6 and TM7 (Fig. 2d-e). In addition, the tail phenyl inserts into the detergent micelle, probably interacts with the lipid bilayer in a native system (Fig. 2f). Compared to peptide bound PTH1R structures, the binding of PCO371 pushes the middle of TM6 outward by ∼8 Å as measured by the Cα of P415^6.47b^ to leave space to accommodate PCO371 (arrows in Extended Fig. 4b&4d).

To investigate the key residues for ligand binding and the receptor activation, we assessed PCO371-induced G_s_ activation by the wild-type and mutant PTH1Rs using cAMP assays. Alanine mutations in hydrophobic pocket residues (I299^3.47b,^ L413^6.45b,^ P415^6.47b,^ and I458^7.56b^) significantly reduced the potency as measured by pEC50 for PCO371 relative to the wild-type PTH1R (Fig.2g, Extended Data Fig. 7b and Extended Data Table 2), indicating these hydrophobic residues play important roles in transmitting PTH1R G-protein signaling. It is in line with the previously reported result that P415^6.47b^ of PTH1R is a key residue for PCO371-mediated PTH1R activation^17^. In addition, alanine substitutions of R219^2.46b^ and Y459^7.57b^ showed clearly a great reduction in the potency of PCO371-mediated G_s_ activation. Alanine substitutions of E302^3.50b^ and H223^2.50b^ also diminished PCO371-induced cAMP production (Fig. 2g, Extended Data Fig. 7b and Extended Data Table 2), which suggests the importance of these residues in PCO371 function. Taken together, the unexpected interface bound by PCO371 between PTH1R and G-protein demonstrates the important roles of individual pocket residues in PCO371 recognition and specificity.

### PTH1R conformational changes and activation

Despite all the active PTH1R structures were solved in the same G protein-bound state^23, 28, 30^, yet they display conformational differences at their TMD bundles between the PCO371-bound and the PTH-bound PTH1R structures. The most notable observation is a 7.5 Å inward shift of at the extracellular end of TM6 in the PCO371-bound PTH1R structure (as measured by the Cα of M425^6.57b,^ Fig. 3a), which causes the extracellular end of TM6 (residues M425^6.57b,^ Y421^6.53b^ and F424^6.56b^) to collide with the PTH N-terminal residues (S1, V2, and S3) (Fig. 3a). This is consistent with the report that PCO371 can inhibit the binding of peptides to their TMD^17^. The conserved PxxG motif (P415^6.47b–^L416^6.48b–^F417^6.49b–^G418^6.50b^) at the middle of TM6 in the PTH-bound PTH1R structure also collide with PCO371 (Fig. 3b), therefore the PxxG motif in the PCO371-PTH1R-G_s_ complex structure is shifted outward to create the binding pocket of PCO371 (Fig. 3b-c). Corresponding to the outward movement of P415^6.47b^ in the PCO371-PTH1R-G_s_ complex structure, the kink of TM6 at P415^6.47b^ is less pronounced than the TM6 kink in the PTH-bound structure (Fig. 3b), leading to less pronounced outward movement (∼4 Å) of TM6 in the cytoplasmic side.

Compared with the PTH-PTH1R-G_s_ complex structure, the PCO371-PTH1R-G_s_ complex structure displays large differences in the extracellular half of the TMD structures but retains very similar structure in the intracellular half of the TMD structure (Extended Data Fig. 4b). Specifically, a large inward movement at the extracellular end of TM6, which pushes large outward movements at the extracellular ends of TM7 and TM1 (Extended Data Fig. 4c). The rearrangement of these structural elements at extracellular side has cascaded into changes of three conserved polar interaction networks in class B GPCR activation as shown in Figure 3d-f. The conformational changes of H420^6.52b^ and Q451^7.49b^ in the central polar network of the PCO371-bound PTH1R structure would resolve the steric clash with F417^6.49b,^ which is flipped upward in the PCO371-bound structure from the PTH-bound structure (Fig. 3d). Y459^7.57b^ from the HETY network is shifted upward and outward to bind with PCO371. The outward shift of P415^6.48b^ resolve the steric clash with conformational changes of Y459^7.57b^ (Fig. 3e). The outward shift of N463^7.61b^ and E465^8.49b^ from the TM2-TM6-TM7-H8 network also resolves the steric clash with each other, and the clash of E465^8.49b^ with R219^2.46b^ (Fig. 3f). These conformational rearrangements together illustrate the structural changes of PTH1R in response to the change of ligand binding from PTH to PCO371, therefore highlighting the capacity of PTH1R to adopt totally different ligands, which induce very distinct receptor conformations in the peptide binding pocket but the receptor can coalesce into a very similar intracellular pocket to couple downstream G proteins.

### The unique aspect of G protein coupling of PTH1R by PCO371

Although the different binding patterns between peptide agonists and the small molecule agonist, PCO371, they activate PTH1R by inducing a consensus kink at the middle of TM6 and subsequent outward shift of the cytoplasmic end of TM6 to form a binding cavity for G protein coupling (Extended Data Fig. 8a-b). Different from the binding modes of all reported peptides and small molecule agonists, PCO371 is at the interface between the receptor and the C-terminus of Gα_s_-α5 in the PCO371-bound PTH1R structure (Extended Data Fig. 8c-e). The C-terminal α5 helix of Gα_s_ makes interactions with TM2, TM3, TM5, TM6 and H8 in both PCO371– and PTH-bound PTH1R structures (Extended Data Fig. 8d-f). In additional, L393 of Gα_s_-α5 forms hydrophobic contact with PCO371, E392 and Y391 of Gα_s_-α5 make polar interactions with PCO371 (Extended Data Fig. 8e). These additional interactions are supported by well-resolved density in the cryo-EM map (Extended Data Fig. 8c) and they can stabilize the active receptor conformation in the G-protein coupling state (R^G^) ^17^. The direct contact of PCO371 with both PTH1R and G protein is consistent with the data reported by Tamura *et al.* ^17^, which has showed that the duration of cAMP response induced by PCO371 is much shorter than that of PTH because PCO371 would bind weakly to PTH1R in the absence of a G protein, consistent with that PCO371 exhibits as an R^G^ –selective ligand ^17^.

### Structural basis of selectivity of PCO371 for PTH1R

To investigate the mechanisms underlying the selectivity of PCO371 for PTH1R over other class B GPCRs, we performed cAMP production assays using transfected wild type receptors of PTH1R, PTH2R and GLP-1R in AD293 cells. PCO371 did not have activity in wild type PTH2R and GLP-1R (Fig. 4a-b). A single residue replacement of L370^6.47b^P of PTH2R converts its response to PCO371-induced activation, while P415^6.47b^A mutation inactivated PTH1R to respond PCO371 but the mutated receptor retained full activation by PTH (Fig. 4a, c). It is worth noting that P^6.47b^ is a conserved residue in TM6 of class B GPCRs except for L370^6.47b^ in PTH2R (Fig. 4d), and our data suggest that P^6.47b^ in PTH receptors is a key residue for the selective activation of PTH receptors by PCO371.

Structure-based sequence alignment of class B GPCRs reveals that the PCO371 binding interface has three non-conserved residues between PTH1R and PTH2R and five non-conserved residues between PTH1R and GLP-1R (Fig. 4d, Extended Data Fig. 9a-e). In contrast to single mutation in PTH2R that can converts its response to PCO371, all single mutations that change GLP-1R residue to PTH1R residue at the five non-conserve PCO371 pocket residues, which mutated receptors retained full activation by GLP-1 peptide, did not convert GLP-1R to respond to PCO371 activation (Extended Data Fig. 9f-g). Combined pocket mutations of two residues, four residues, or five residues can convert the mutated GLP-1R to be activated by PCO371 but not by PTH (Fig. 4b-c). The degree of PCO371 activation by the two-residue mutated GLP-1R is the same (if not better) as that by the four-residue or five-residue mutated GLP-1R, suggesting these two residues are key for PCO371 selectivity.

### A conserved binding site in class B GPCRs for small molecule ligands

The ability of PCO371 activation by one-residue mutated PTH2R or two-residue mutated GLP-1R suggest a possibility of a similar PCO371 binding pocket conserved in members of class B GPCRs. To validate this hypothesis, we performed sequence alignment and homology modeling based on the PCO371-bound PTH1R structure (Fig. 4d and Fig. 5a). Sequence alignment reveals that most residues of the 15 PTH1R residues that comprise the PCO371 pocket are conserved across class B GPCRs (Fig. 4d). Structural modeling of all other members of class B GPCRs suggest the existence of a similar PCO371 binding pocket in these receptors, in which PCO371 could adopt a similar binding mode to the PCO371-PTH1R structure (Fig. 5a). To corroborate the sequence and structure analyses, we tested the ability of PCO371 to activate other members of class B GPCRs (Fig. 5b). In addition to PTH1R, seven wildtype class B GPCRs (GCGR, GIPR, PAC1R, GHRHR, SCTR, VIP1R, and VIP2R) can be activated by PCO371 (Fig 5b). For GLP-1R, GLP-2R, PTH2R, and CRF2R, their wildtype receptors cannot be activated by PCO371 but one or two mutations in the pocket residues can convert them to respond to PCO371 activation. Based on these results, we conclude that a PCO371-like pocket is mostly conserved in class B GPCRs.

## Conclusions

In summary, the structure of PCO371-bound PTH1R-G_s_ complex provides a structural basis of small molecule agonist binding and activation of PTH1R. This work reveals an unanticipated small molecule agonist-binding site and serve as a template for homology modelling of class B GPCRs. The PCO371 binding site is within the TMD at the interface with G protein, which is far away from the receptor ECD, thus small molecule agonists at this site may not require to mimic the interactions of peptides with ECD to promote the binding affinity. Class B GPCRs have higher sequence homology in their TMDs than their ECDs. Our modeling and receptor activation studies suggest that a PCO371-like pocket is likely conserved in most members of class B GPCRs, thus providing a general and exciting direction for structure-based design of small-molecule drugs targeting this new binding site at class B GPCRs.

## Materials and Methods

### Constructs of PTH1R and heterotrimeric G proteins

The human PTH1R (residues 27-502) with G188A and K484R mutations was cloned into pFastBac vector (Invitrogen) with the haemagglutinin signal peptide (HA), followed by a TEV protease cleavage site and a double MBP (2MBP) and His tag to facilitate expression and purification^28^. To facilitate a stable complex, the above PTH1R construct was added the LgBiT subunit (Promega) at the C terminus of PTH1R with a 17-amino acid linker. Based on the published DNGα_s_, a modified bovine Gα_s_ (mDNGα_s_), its N terminus (M1–K25) and α-helical domain (AHD F68–L203) of Gα_s_ were replaced with the N terminus (M1–M18) and AHD (Y61–K180) of the human Gα_i_, which can bind scFv16 and Fab_G50^31^ and the residues N254-T263 of Gα_s_ were deleted. In addition, eight mutations (G49D, E50N, L63Y, A249D, S252D, L272D, I372A, and V375I) were added to improve stability of G protein subunits^32^. To facilitate the folding of the G protein, mDNGα_s_ was co-expressed with GST-Ric-8B^33^. Rat Gβ_1_ was fused with a His-tag at the N terminus and with a SmBiT subunit (peptide 86, Promega)^34^ after a 15-amino acid linker at its C terminus. The wild type (WT) and mutants of PTH1R, PTH2R, GLP-1R, GLP-2R, GCGR, GIPR, GHRHR, SCTR, PAC1R, VIP1R,

VIP2R and CRF2R were constructed into the pcDNA6.0 vector (Promega) for cAMP accumulation. PTH1R, β-arrestin1 and β-arrestin2 were constructed into pBiT vector for arrestin recruitment. All constructs were cloned using Phanta Max Super-Fidelity DNA Polymerase (Vazyme Biotech Co., Ltd).

### Expression of PCO371**-**PTH1R**-**G_S_ complex

To facilitate a stable complex assembly and purification, PTH1R and G proteins were co-expressed in *Sf9* insect cells (Invitrogen). The *Sf9* cells grew to a density of 3.5 × 10^6^ cells/mL in ESF 921 cell culture medium (Expression Systems) for expression. We infected the cells with five separate virus preparations at a ratio of 1:2:2:2:2, including PTH1R (27-502)-17AA-LgBiT-2MBP, mDNGα_s_, Gβ_1_-peptide 86, Gγ_2_, and GST-Ric-8B. The infected cells were cultured at 27°C for 48 h, the cells were harvested by centrifugation and washed with PBS once. The cell pellets were frozen at −80°C for further usage.

### Expression and purification of Nb35

Nanobody-35 (Nb35) was expressed in *E. coli* BL21 cells, the cultured cells were grown in 2TB media with 100 μg/mL ampicillin, 2 mM MgCl_2_, 0.1% glucose at 37°C for 2.5 h until OD600 of 0.7-1.2 was reached. Then the culture was induced with 1 mM IPTG at 37°C for 4-5 h, and harvested and frozen at –80°C for further purification. Nb35 was purified by nickel affinity chromatography and followed by size-exclusion chromatography using HiLoad 16/600 Superdex 75 column or following overnight dialysis against 20 mM HEPES, pH 7.4, 100 mM NaCl, 10% glycerol. The Nb35 protein was verified by SDS-PAGE and store at −80 °C.

### Purification of PCO371**-**PTH1R**-**G_S_ complex

The complex was purified according to previously described methods^28, 35^. The cell pellets were resuspended in 20 mM HEPES pH 7.4, 100 mM NaCl, 10 mM MgCl_2_, 10 mM CaCl_2_, 2 mM MnCl_2_, 10% glycerol, 0.1 mM TCEP, 15 μg/mL Nb35, 25 mU/mL apyrase (Sigma), 200 µM PCO371 (Hefei Fuya Biotechnology Co., Ltd), supplemented with Protease Inhibitor Cocktail (TargetMol, 1 mL/100 mL suspension). The lysate was incubated for 1 h at room temperature and then solubilized by 0.5% (w/v) lauryl maltose neopentylglycol (LMNG, Anatrace) supplemented with 0.1% (w/v) cholesteryl hemisuccinate TRIS salt (CHS, Anatrace) for 2 h at 4°C. The supernatant of the solubilized membranes was collected by centrifugation at 65,000 × g for 40 min, then incubated with Amylose resin (Smart-lifesciences) for 2 h at 4°C. The resin was loaded onto a gravity flow column and washed with 20 column volumes of 20 mM HEPES, pH 7.4, 100 mM NaCl,10% glycerol, 5 mM CaCl_2_, 5 mM MgCl_2_, 1 mM MnCl_2_, 0.01% (w/v) LMNG, 0.01% glyco-diosgenin (GDN, Anatrace) and 0.004% (w/v) CHS, 100 µM PCO371, and 25 μM TCEP. After washing, the protein was cut with TEV protease on column overnight at 4°C. Next day the flow through was collected and concentrated, then PCO371-PTH1R-G_s_ flow through was loaded onto a Superdex200 10/300 GL column (GE Healthcare), with the buffer consisting of 20 mM HEPES, pH 7.4, 100 mM NaCl, 2 mM MgCl_2_, 0.00075% (w/v) LMNG, 0.00025% GDN, 0.0005% (w/v) digitonin (Biosynth), 0.0002% (w/v) CHS, 50 µM PCO371, and 100 μM TCEP. The complex fractions were collected and concentrated for electron microscopy experiments.

### Cryo-EM grid preparation and data acquisition

For cryo-EM grid preparation of PCO371-PTH1R-Gs complex, 3.0 μL purified protein at a concentration of ∼4.95 mg/mL was used for the glow-discharged holey carbon grids (Quantifoil, R1.2/1.3, Au, 300 mesh). The grids were blotted for 2s at 4°C, in 100% humidity using a Vitrobot Mark IV (Thermo Fisher Scientific) and then plunge-frozen in liquid ethane. The frozen grid of PCO371-PTH1R-G_s_ complex was transferred to a Titan Krios G4 equipped with a Gatan K3 direct electron detector and cryo-EM movies were performed automatic data collection. with super-resolution mode at a pixel size of 0.412 Å using EPU at Advanced Center for Electron Microscopy at Shanghai Institute of Materia Medica, Chinese Academy of Sciences. A total of 8,002 Movies were recorded with pixel size of 0.824 Å at a dose of 50 electron per Å^2^ for 36 frames. The defocus range of this dataset was –0.8 µm to –1.8 µm. For dimer complex, another 5,364 movies ware obtained with same parameters.

### Cryo-EM data processing

All dose-fractionated image stacks were subjected to beam-induced motion correction by Relion 4.0^36^. The defocus parameters were estimated by CTFFIND 4.1^37^ of Cryosparc^38^. For PCO371-PTH1R-G_s_ dataset, template auto-picking yielded 7,124,33 particles, which were processed two rounds by reference-free 2D classification using Cryosparc^38^. With initial model, after two rounds of 3D classification using Relion, local masks were used on receptor. 1,099,315 particles were used to further refinement and polishing. Particle subtractions were used on complex to subtract micelle and do refinement, yielding reconstructions with global resolution of 2.57 Å, and subsequently post-processed by DeepEMhancer^39^.

### Model building and refinement

The cryo-EM structure of the LA-PTH1R-G_s_-Nb35 complex (PDB code 6NBF) was used as the start for model building and refinement against the electron microscopy map. The model was docked into the electron microscopy density map using Chimera^40^, followed by iterative manual adjustment and rebuilding in COOT^41^. Real space and Rosetta refinements were performed using Phenix^42^. The model statistics were validated using MolProbity^43^. Fitting of the refined model to the final map was analyzed using model-versus-map FSC. To monitor the potential over-fitting in model building, FSC_work_ and FSC_free_ were determined by refining ‘shaken’ models against unfiltered half-map-1 and calculating the FSC of the refined models against unfiltered half-map-1 and half-map-2. The final refinement statistics are provided in Supplementary Table 2. Structural figures were prepared in Chimera and PyMOL (https://pymol.org/2/).

### Modeling and volume calculation

The homology modeling of class B GPCRs was based on the PTHR structure using MODELLER^44^. The sequence of PTHR in our cryo-EM structure was used as the reference sequence. After alignment from the receptor sequence from other class B GPCR structures, AutoModel of MODELLER was applied for homology modeling. The structure with the lowest Discrete Optimized Protein Energy (DOPE) potential was used for the following volume calculation using PyVOL^45^. In volume calculation, the minimum radius was 1.2, while the maximum radius was 3.4. The pocket was defined as the residues around 5 Å of ligand and during calculation, the Gα protein of PTHR was kept.

### cAMP accumulation assay

PTH, PCO371, TIP39 and GLP-1 stimulated cAMP accumulations were measured by a LANCE Ultra cAMP kit (PerkinElmer). After 24 h culture, the transfected AD293 cells were seeded into 384-well microtiter plates at a density of 3,000 cells per well in HBSS supplemented with 5 mM HEPES, 0.1% (w/v) BSA or 0.1% (w/v) casein and 0.5 mM 3-isobutyl-1-methylxanthine. The cells were stimulated with different concentrations of peptide agonists for 30 min at RT. Eu-cAMP tracer and ULight^TM^– anti-cAMP were then diluted by cAMP detection buffer and added to the plates separately to terminate the reaction. Plates were incubated at RT for 15min and the fluorescence intensity measured at 620 nm and 665 nm by an EnVision multilabel plate reader (PerkinElmer).

### NanoBiT β-Arrestin recruitment assay

The recruitment of PTH1R to β-arrestin was detected in HEK293 cells using the NanoLuc Binary System (NanoBiT; Promega). The Lgbit subunit was fused to the C-terminus of PTH1R and the SmBiT subunit was fused to the N-terminus of β-arrestin. The HEK293 cells were harvested and plated into 384-wells microtiter plates at a density of 3000 cells per well 24 h after co-transfection of PTH1R-LgBiT and SmBiT– β-arrestin. Coelenterazine was then added to the plates in the dark with the final concentration of 10 μM (5μL/well). The ligands of different concentrations were finally added to the plates and then the bioluminescence signal was measured using an EnVision plate reader (PerkinElmer).

### Surface expression assay

Surface expression of PTH1R WT and mutants were cloned into pcDNA6.0 (Invitrogen) with 3× Flag tag at C-terminal and determined by flow cytometry. AD293 cells were collected after 24 hours of transient transfection and then blocked with 5% BSA in PBS at RT for 15 min followed by incubation with primary mouse anti-Flag antibody at RT for 1 hour. The cells were then washed three times with PBS containing 1% BSA and incubated with anti-mouse Alexa-488-conjugated secondary antibody at 4 ℃ in the dark for 1h. After another three washes, the cells were resuspended with 500 μl PBS containing 1% BSA for detection in BD Accuri C6 flow cytometer system (BD Biosciences) at excitation 488 nm and emission 519 nm. For each sample, approximately 5000 cellular events were collected and the data were normalized to PTH1R WT.

### Statistical analysis

All functional data were displayed as means ± standard error of the mean (S.E.M.). Statistical analysis was performed using GraphPad Prism 8.0 (GraphPad Software). Experimental data were evaluated with a three-parameter logistic equation. The significance was determined with either two-tailed Student’s t-test or one-way ANOVA. *P* < 0.05 was considered statistically significant.

## Data availability

Cryo-EM map has been deposited in the Electron Microscopy Data Bank under accession code: EMD-XXXX (PCO371-bound PTH1R-G_s_ complex). The atomic coordinate has been deposited in the Protein Data Bank under accession codes: XXXX (PCO371-bound PTH1R-G_s_ complex).

**Extended Data Fig. 1.**
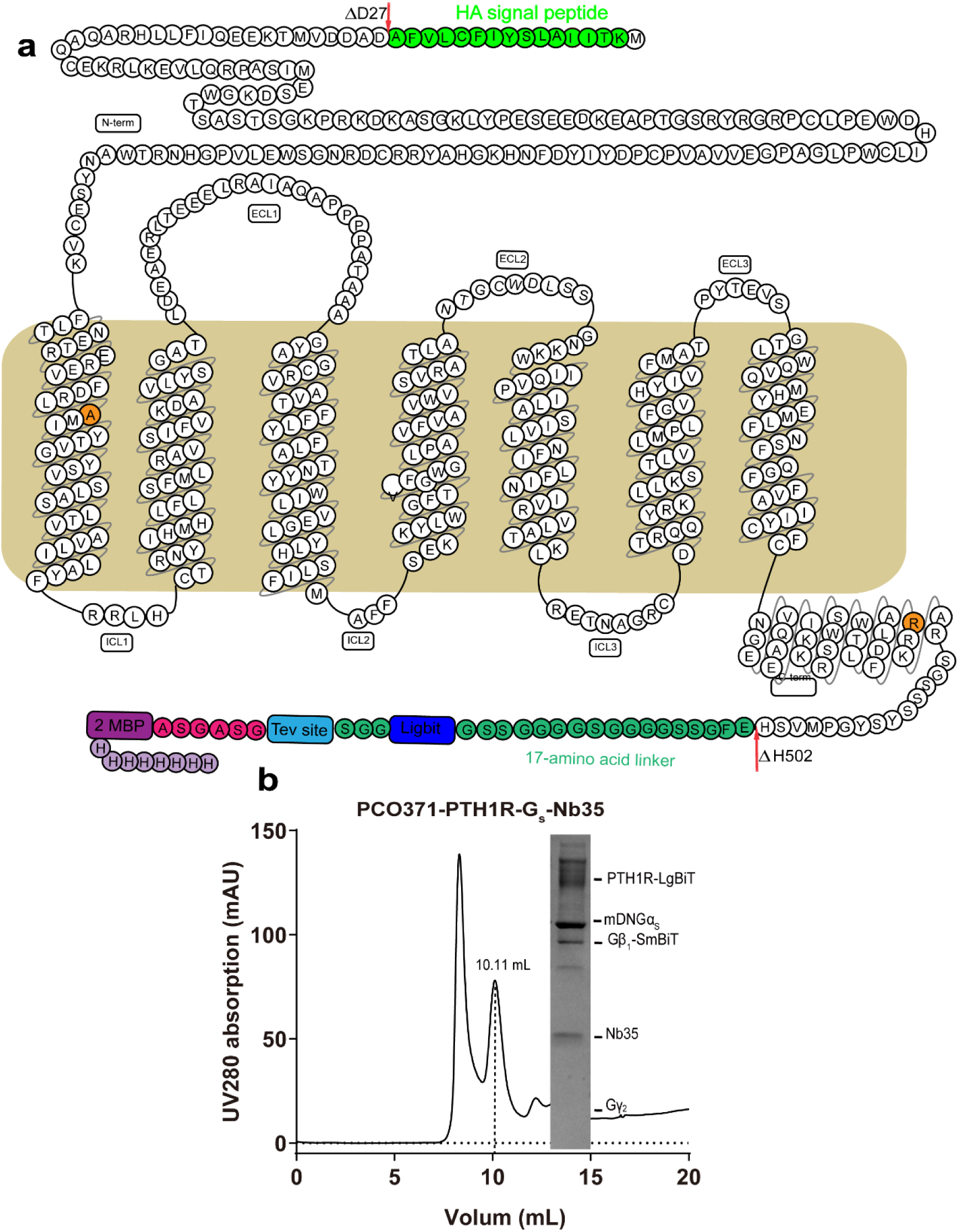
| Construct of receptor and purification of the PCO371-PTH1R-G_s_ complex. (**a**) Snake plot diagram of the PTH1R-LgBiT construct. (**b**) The size-exclusion chromatography elution profile on Superdex200 Increase 10/300GL (left panel) and SDS-PAGE analysis (right panel) of the PCO371-PTH1R-G_s_ complex.

**Extended Data Fig. 2.**
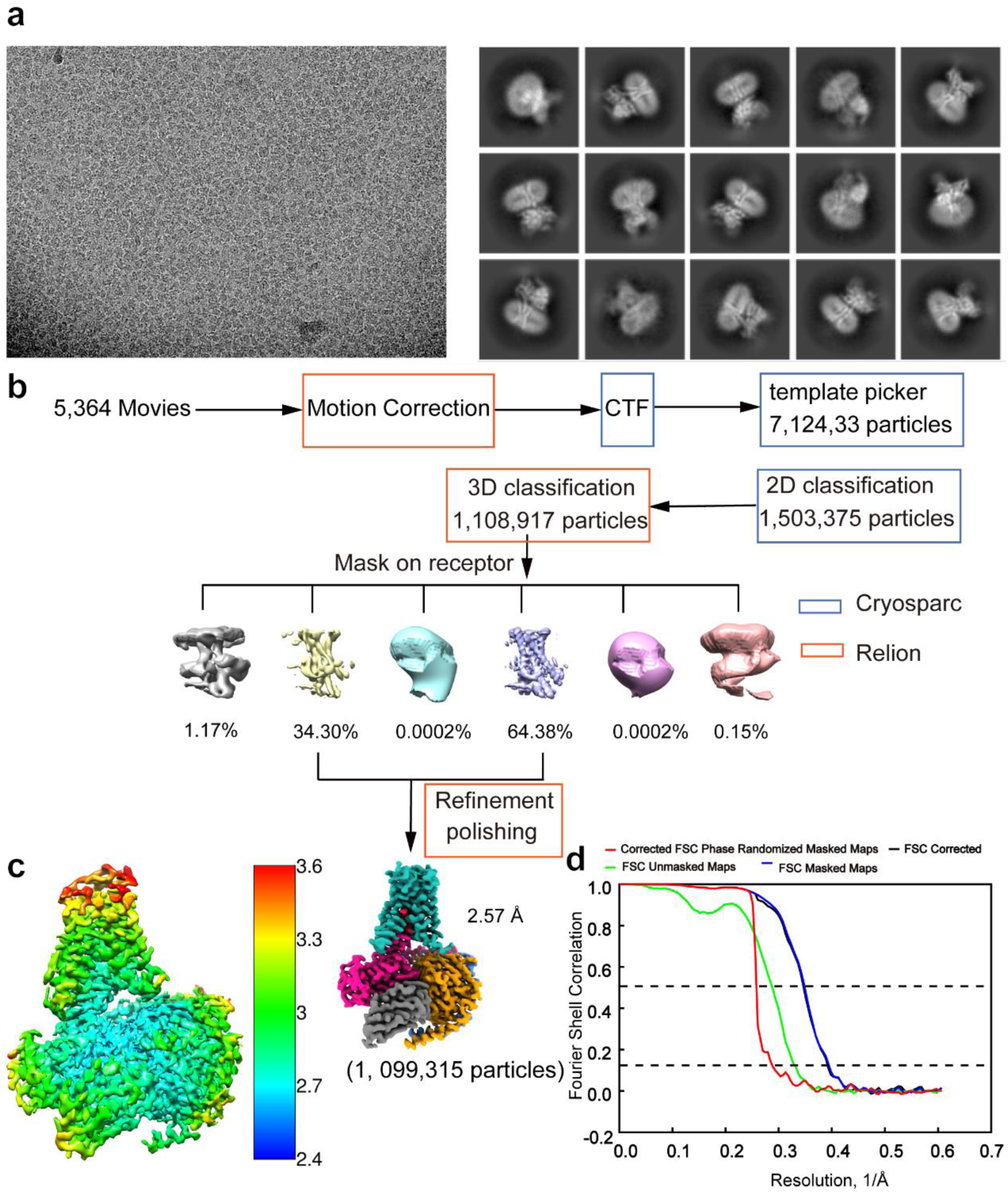
| Single particle cryo-EM data analysis of the PCO371-PTH1R-G_s_ complex. (**a**) A representative cryo-EM micrograph of the PCO371-PTH1R-G_s_ complex and representative 2D class averages with distinct secondary structure features from different views. (**b**) Data processing flowchart of PCO371-PTH1R-G_s_ complex by CryoSPARC and Relion. (**c**) Color cryo-EM map of the PCO371-PTH1R-G_s_ complex, showing local resolution (Å) calculated using Relion. (**d**) “Gold-standard” FSC curve of the PCO371-PTH1R-G_s_ complex, with the global resolution defined at the FSC = 0.143 is 2.57 Å.

**Extended Data Fig. 3.**
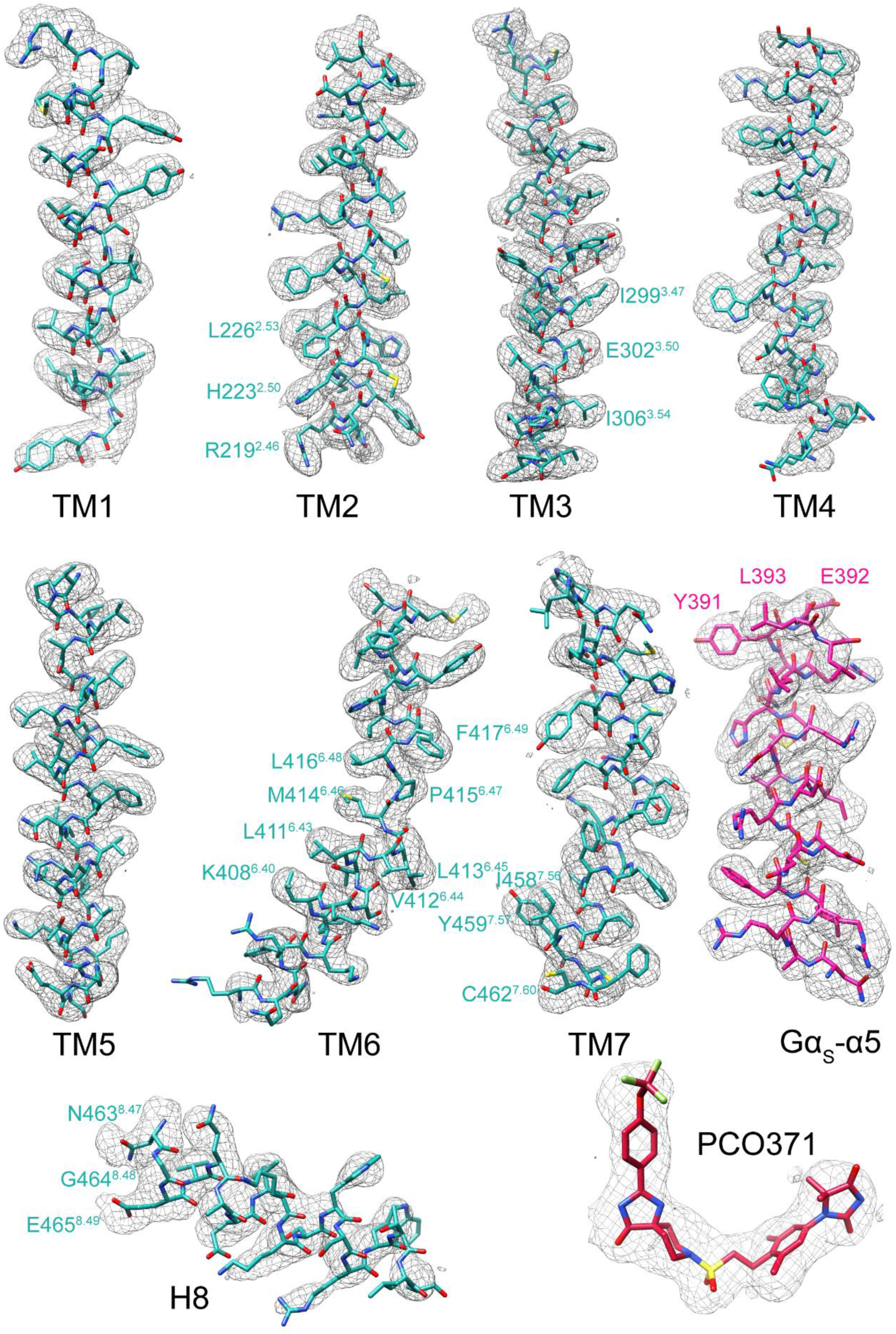
| Cryo-EM density maps of the PCO371-PTH1R-G_s_ protein structures. Cryo-EM density map and the model of the PCO371-PTH1R-G_s_ structure are shown for all transmembrane helices and helix 8 of PTH1R, PCO371, and Gα_s_-α5 helix. The model is shown in stick representation. All of them display good density.

**Extended Data Fig. 4.**
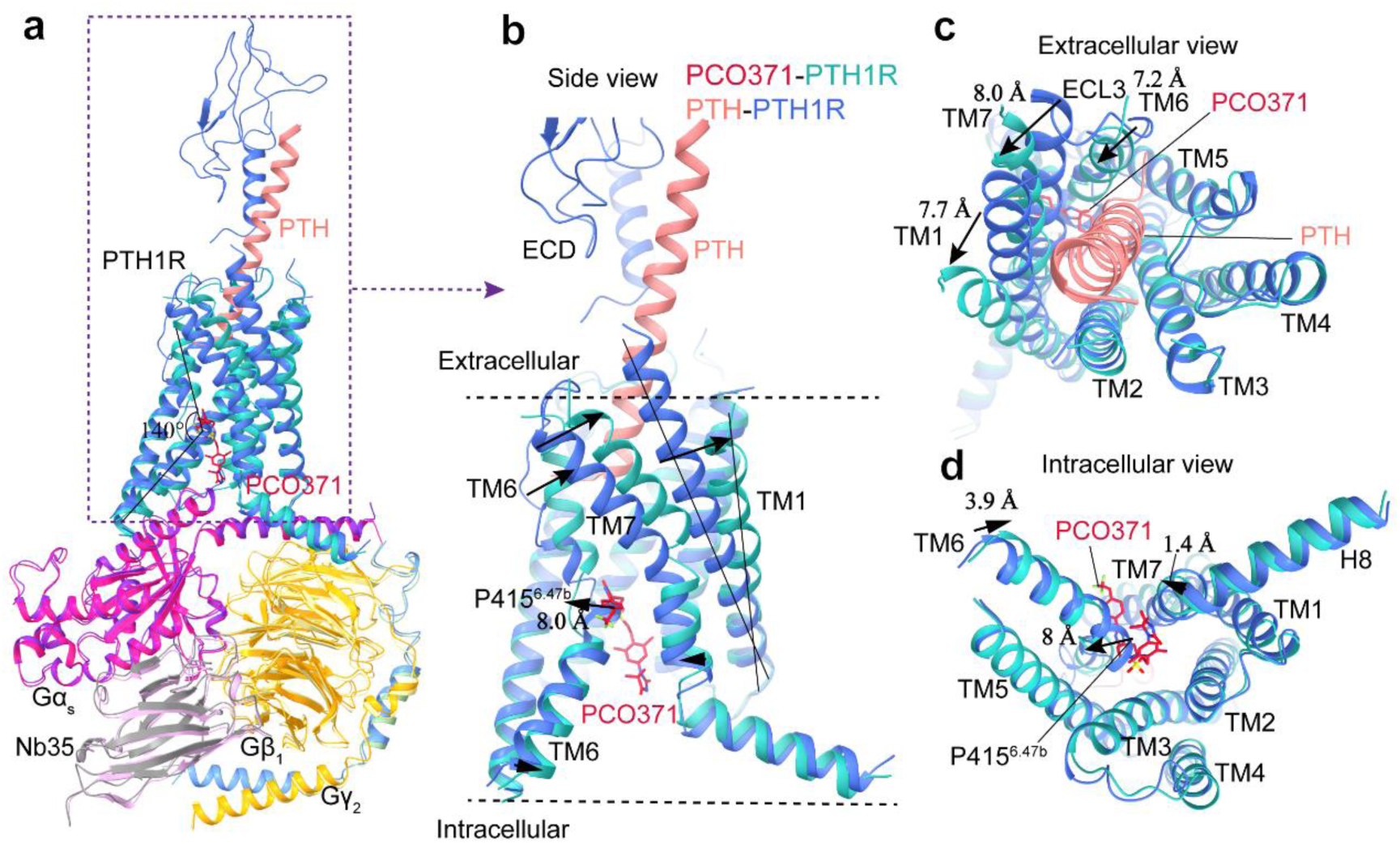
| Comparisons of the agonist binding pockets and PTH1R conformations stabilized by PCO371 and PTH. (**a**) Superimposition of PTH1R from PDB:8HA0 (PTH1R: royal blue, PTH34: light coral, Gα_s_: blue violet, Gβ_1_: khaki, Gγ_2_: dark sea green, Nb35: plum) and the PCO371-bound PTH1R structure (PTH1R: light sea green, PCO371: crimson, Gα_s_: deep pink, Gβ_1_: orange, Gγ_2_: cornflower blue, Nb35: gray) reveals different peptide– and PCO371-binding sites. (**b**) Side view of different binding pockets and conformational changes in receptors; (**c**) Extracellular view and (**d**) intracellular view of PTH1R conformational changes.

**Extended Data Fig. 5.**
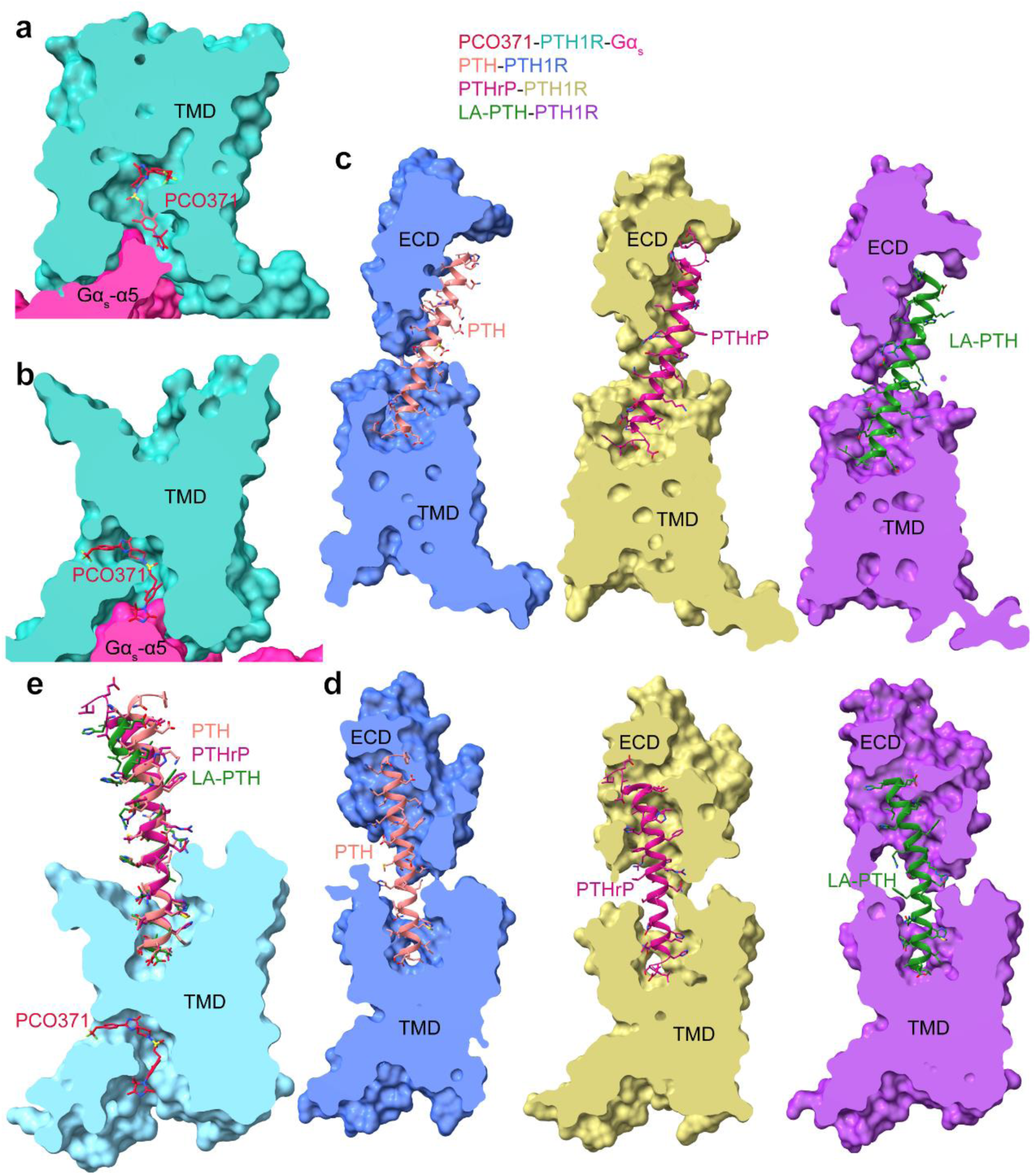
| Differences of ligand-binding pockets between a small molecule agonist and peptides of PTH1R. (**a-b**) The binding pocket of PCO371 in PCO371-PTH1R-G_s_ complex structure. The receptor is shown in surface representation and colored in light sea green and PCO371 in crimson is shown as sticks. G protein and Nb35 are omitted for clarity. (**c-d**) The binding pockets of different peptides of PTH1R in the G protein-bound state. In three PTH-, PTHrP– and LA-PTH-bound PTH1R-G_s_ complex structures, the receptors are shown in surface representation and colored in royal blue, dark khaki and dark orchid, respectively. PTH, PTHrP and LA-PTH are colored in light coral, medium violet red and forest green, respectively. They are shown as sticks and ribbon (PDB: 8HA0, 8HAF and 6NBF). G protein and Nb35 are omitted for clarity.

**Extended Data Fig. 6.**
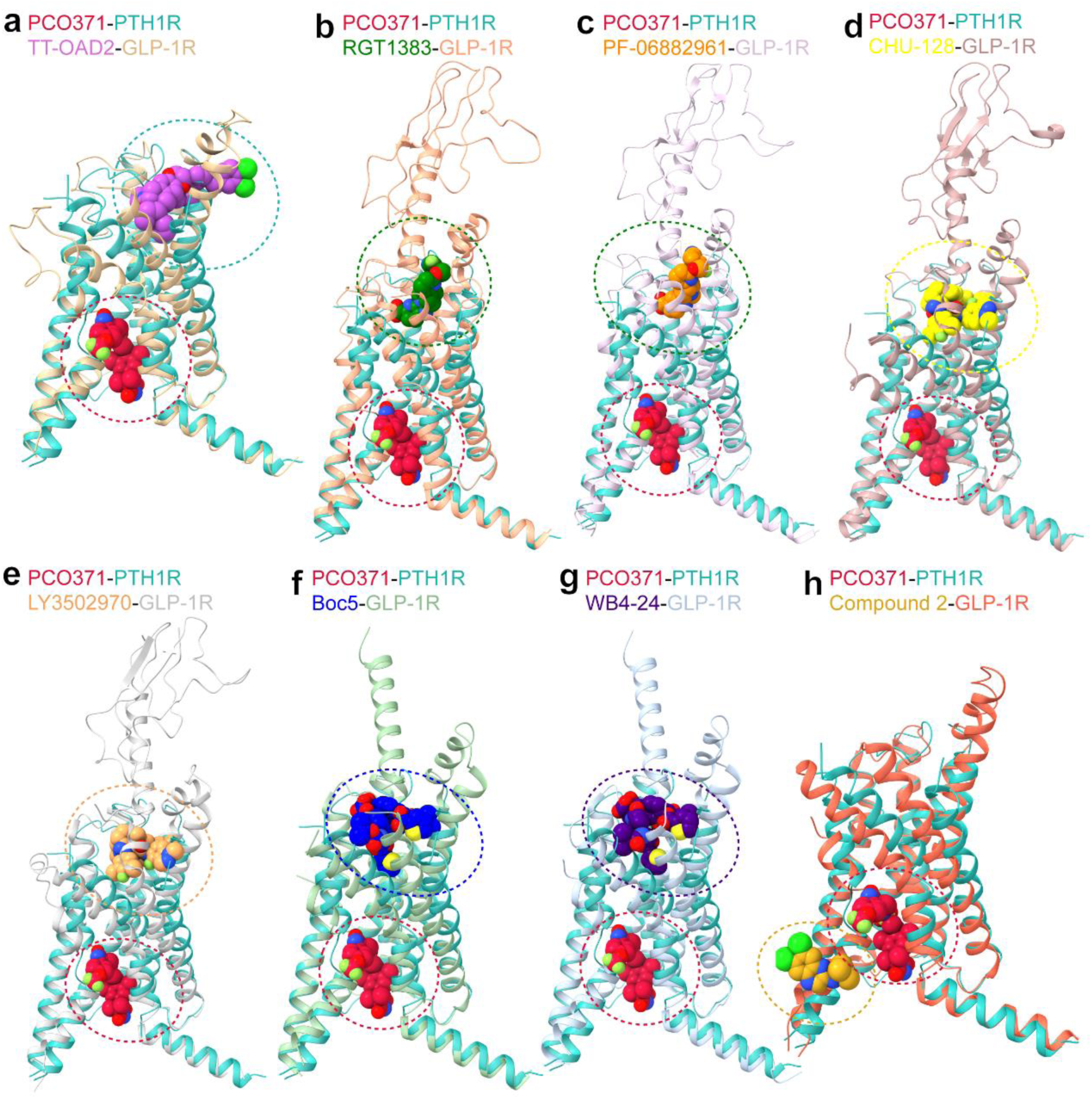
| Comparisons of small molecule agonist binding sites of class B GPCRs. (**a-h**) Comparisons of the overall backbone conformations of helical bundles and the ligand binding pockets between PCO371-PTH1R-G_s_ and non-peptidic ligand– GLP-1R-G_s_ complexes. Superimposition of the PTH1R (light sea green) in complex with G_s_ bound to PCO371 (crimson) with the GLP-1R in complexes with G_s_ bound to different non-peptidic ligands, including small molecule agonists: TT-OAD2(PDB: 6ORV; TT-OAD2: dark orchid, GLP-1R: burly wood); RGT1383 (PDB: 7C2E; RGT1383: green, GLP-1R: light salmon); PF-06882961(PDB: 6X1A; PF-06882961: dark orange, GLP-1R: thistle); CHU-128 (PDB: 6X19; CHU-128: yellow, GLP-1R: rosy brown); LY3502970 (PDB: 6XOX; LY3502970: sandy brown, GLP-1R: silver); Boc5 (PDB: 7×8r; Boc5:blue, GLP-1R: dark sea green) and WB4-24 (PDB:7×8s; WB4-24: indigo, GLP-1R: light steel blue) and with an allosteric ligand, Compound 2, (PDB: 7EVM; Compound 2: goldenrod, GLP-1R: tomato). Gα_s_, Gβ_1_ and Gγ_2_ were omitted for clarity. (**a**) PCO371-PTH1R-G_s_ and TT-OAD2-GLP-1R-G_s_ complexes. (**b**) PCO371– PTH1R-G_s_ and RGT1383-GLP-1R-G_s_ complexes. (**c**) PCO371-PTH1R-G_s_ and GLP-1R-PF-06882961-G_s_ complexes. (**d**) PCO371-PTH1R –G_s_ and CHU-128-GLP-1R-G_s_ complexes. (**e**) PCO371-PTH1R-G_s_ and LY3502970-GLP-1R-G_s_ complexes. (**f**) PCO371-PTH1R-G_s_ and Boc5-GLP-1R-G_s_ complexes. (**g**) PCO371-PTH1R-G_s_ and WB4-24-GLP-1R-G_s_ complexes. (**h**) PCO371-PTH1R-G_s_ and Compound 2-GLP-1R-G_s_ complexes. G protein and Nb35 are omitted for clarity.

**Extended Data Fig. 7.**
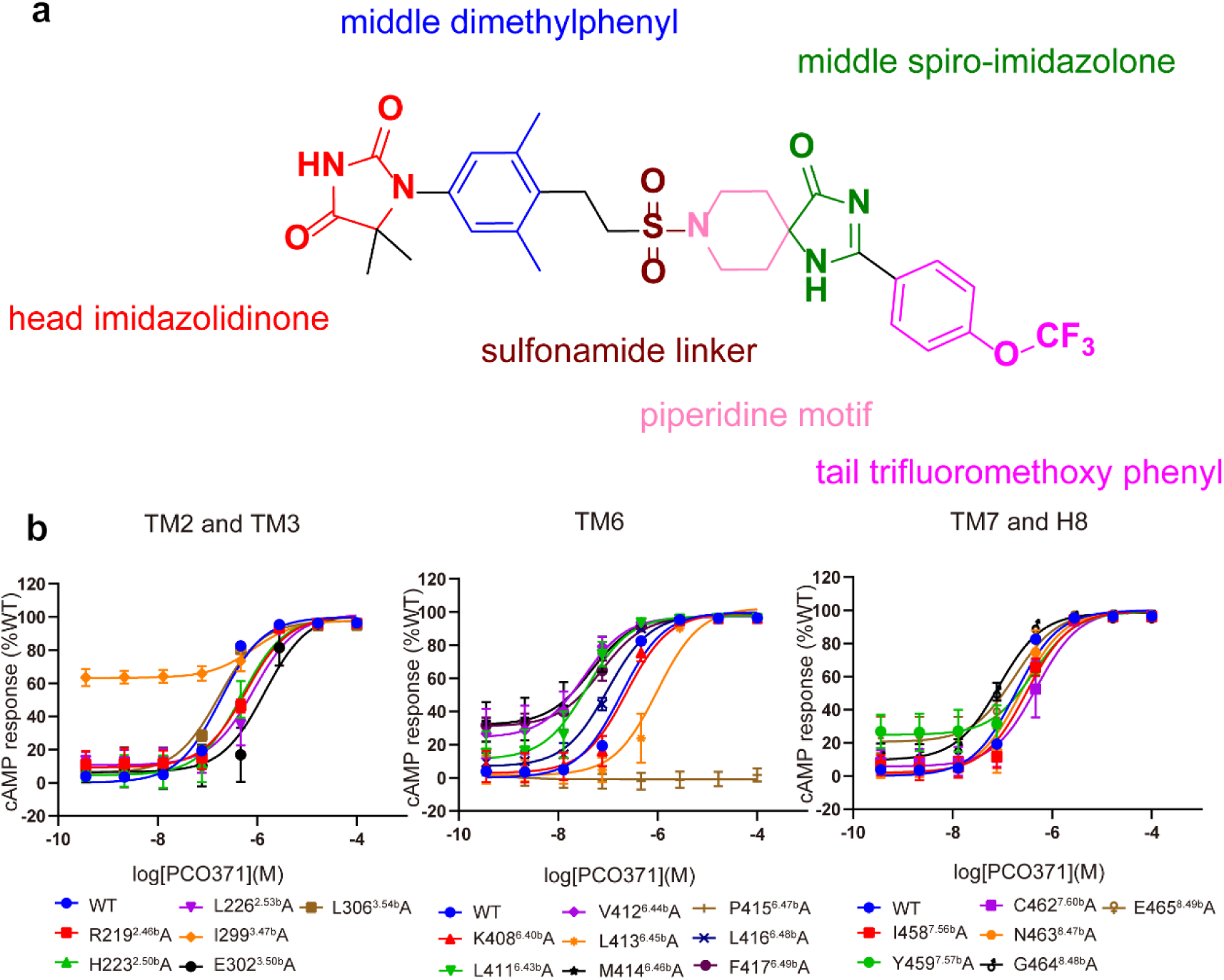
| Chemical structure of PCO371 and PCO371-mediated cAMP production by receptors containing alanine mutants of key residues in PCO371 binding pocket. (**a**) The chemical structure of PCO371 is comprised of the head imidazolidinone, the middle dimethylphenyl, the sulfonamide linker, the piperidine motif, the middle spiro-imidazolone, and the tail trifluoromethoxy phenyl. (**b**) PCO371-mediated cAMP production by receptors containing alanine mutants of key residues within TM2, TM3, TM6, TM7 and H8. Data from three independent experiments (n=3) performed in technical triplicate are presented as mean ± SEM.

**Extended Data Fig. 8.**
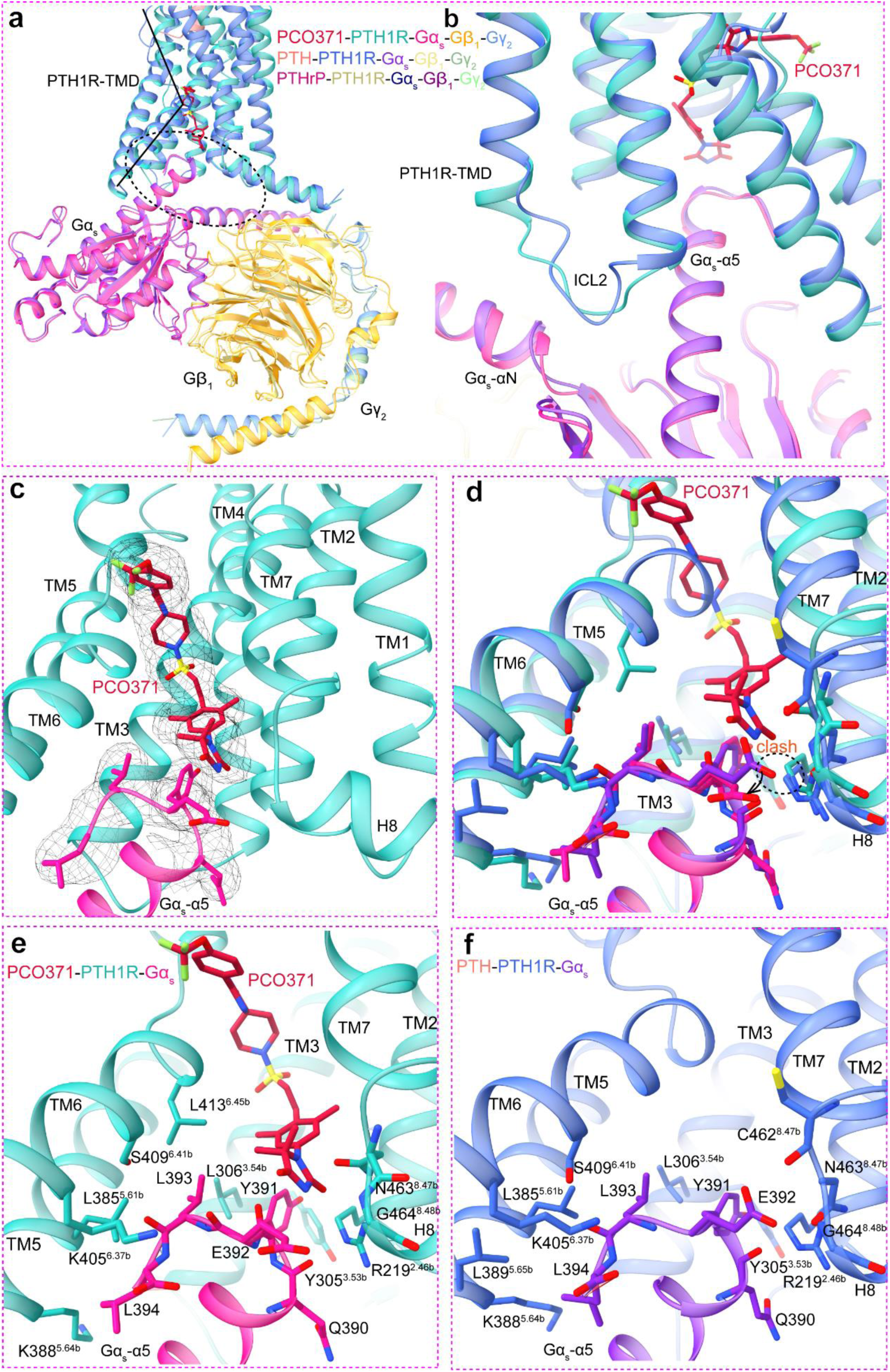
| The similarity and the difference of PTH1R in G protein-coupling by hormone peptide and small molecule agonist. (**a**) Structural comparison of G protein in different ligands bound PTH1R-G_s_ complex structures. (**b**) Close up of the αN and Gα_s_-α5 helix of Gα_s_, which form interactions with ICL2 and TMD helix bundles in all G protein bound complex structures, showing similar G protein conformation, but the noteworthy difference is that the C-terminal of Gα_s_-α5 helix makes additional interactions with the small molecule agonist. (**c**) Good cryo-EM density supports ligand interact with Gα_s_. (**d-f**) The similar set of interactions between the C-terminal of Gα_s_-α5 helix with the receptor. E392 shifts outward due to steric clash. Y391, E392, and L393 form additional interactions with PCO371.

**Extended Data Fig. 9.**
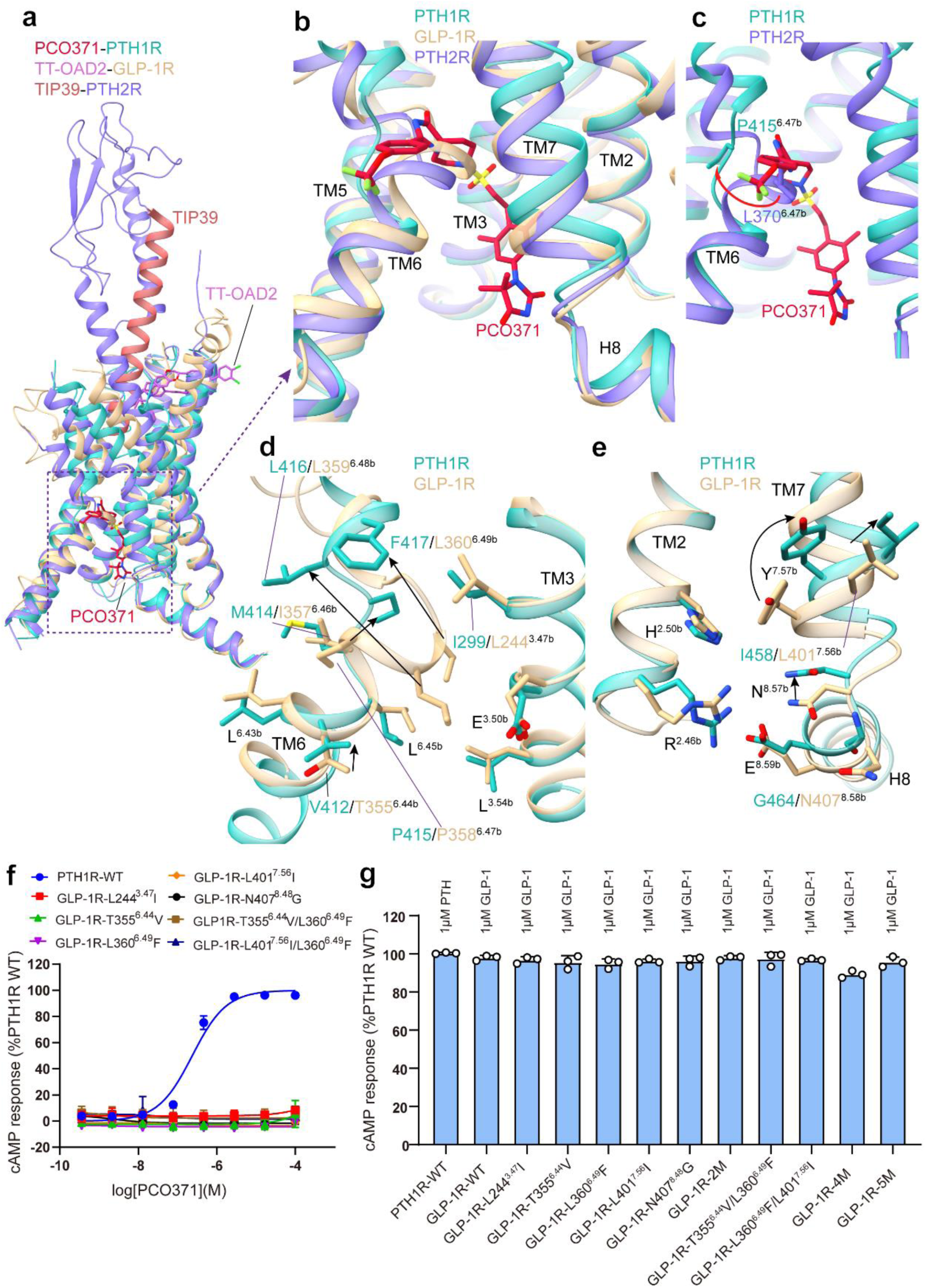
| Key residues for PCO371 selectivity among class B GPCRs. (**a**) Structural comparison of receptors and ligands among PCO371-PTH1R-Gs and TT-OAD2-GLP-1R-Gs and TIP39-PTH1R-Gs complexes. (**a-b**) Structural comparison of the cytoplasmic regions of PTH1R, PTH2R and GLP-1R during the receptor activation. (**c**) Structural comparison of P415^6.47b^ and L370^6.47b^ in PTH receptors. (**d-e**) Different conformations of residues in the active PTH1R, and GLP-1R that are involved the interface of PCO371 in receptor activation. (**f**) Stimulation of cAMP production by PCO371 in the WT and mutants of GLP-1R. Data from three independent experiments (n=3) performed in technical triplicate are presented as mean ± SEM. (g) Stimulation of cAMP production by the cognate ligands of PTH1R, PTH2R and GLP-1R in mutants of receptors. Data from three independent experiments (n=3) performed in technical triplicate are presented as mean ± SEM.

**Extended Data Table 1.** | Cryo-EM data collection, refinement and validation statistics.

|  |  |
| --- | --- |
|  | PCO371-PTH1R-G<sub>s</sub>-complex |
| **Data collection and processing** |  |
| Magnification | 105000 |
| Voltage (kV) | 300 |
| Electron exposure (e<sup>-</sup>/Å<sup>2</sup>) | 50 |
| Defocus range (μm) | -1.2 to -1.8 |
| Pixel size (Å) | 0.824 |
| Symmetry imposed | C1 |
| Initial particle images (no.) | 7,124,33 |
| Final particle images (no.) | 1,099,315 |
| Map resolution (Å) |  |
| FSC threshold | 0.143 |
| Map resolution (Å) | 2.57 |
| Map sharpening B factor (Å<sup>2</sup>) | -69.24 |
| Refinement |  |
| Initial model used (PDB code) | 6NBF |
| Model resolution (Å) | 3.1 |
| FSC threshold | 0.5 |
| Model-Map CC (mask) | 0.81 |
| Model composition |  |
| Non-hydrogen atoms | 8138 |
| Protein residues | 1019 |
| B factors (Å<sup>2</sup>) |  |
| Protein | 66.78 |
| Ligand | 60.37 |
| R.m.s. deviations |  |
| Bond lengths (Å) | 0.002 |
| Bond angles (Å) | 0.527 |
| Validation |  |
| MolProbity score | 1.40 |
| Clash score | 7.2 |
| Rotamer outliers (%) | 0 |
| Ramachandran plot |  |
| Favored (%) | 98 |
| Allowed (%) | 2 |
| Disallowed (%) | 0 |

**Extended Data Table 2.**
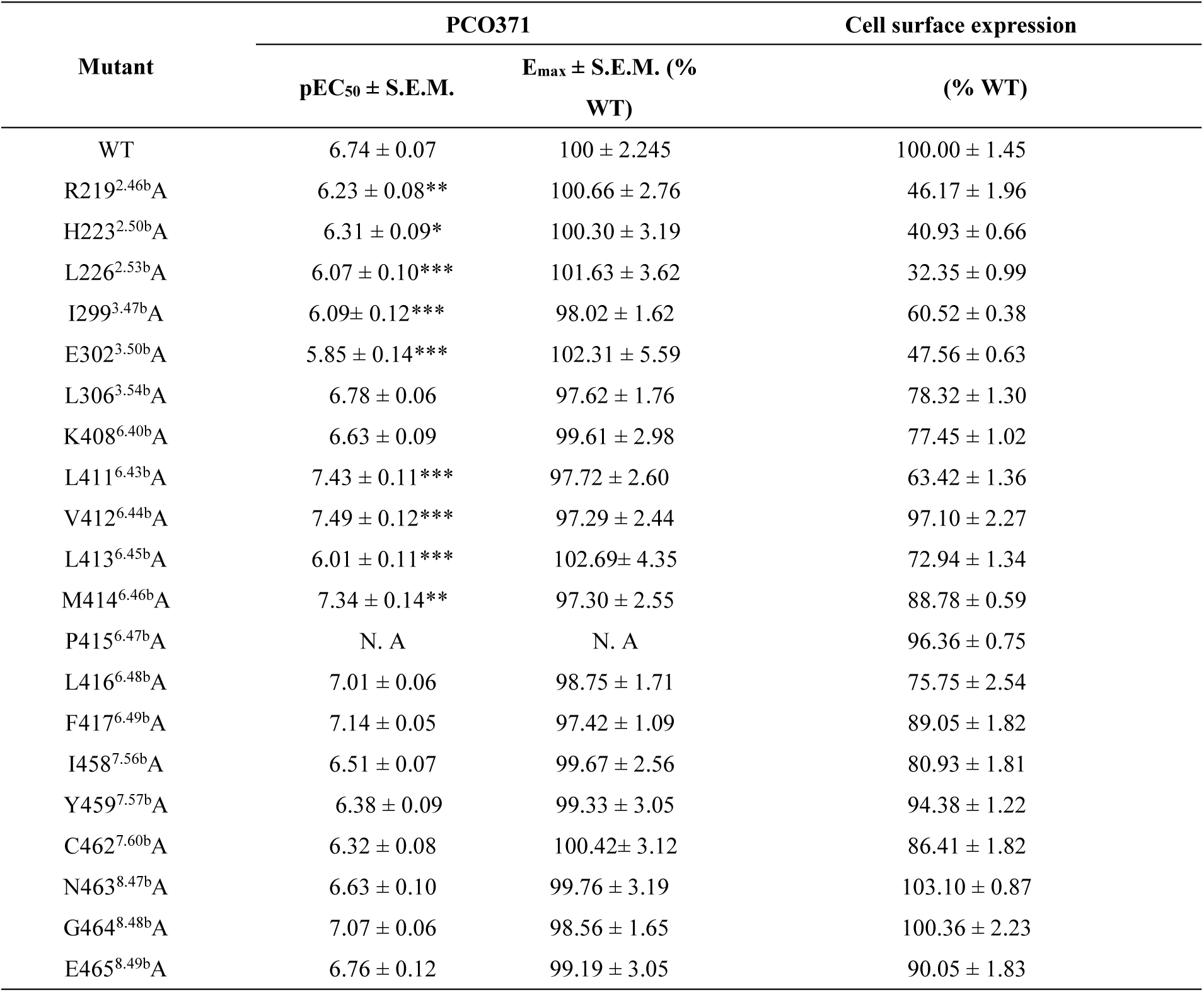
| Effects of PCO371 bind to PTH1R WT and mutants.

## Acknowledgements

The cryo-EM data were collected at Advanced Center for Electron Microscopy at Shanghai Institute of Materia Medica, Chinese Academy of Sciences. We are grateful to Wen Hu and Kai Wu for collecting the cryo-EM data. This work was supported by the Young Innovator Association of CAS (2018325 to LHZ); National Natural Science Foundation of China (32071203 to LHZ, 32130022 and 82121005 to H.E.X.); the National Key R&D Program of China (2019YFA0904200) and SA-SIBS Scholarship Program to LHZ; Ministry of Science and Technology (China) grants (2018YFA0507002 to H.E.X.); Shanghai Municipal Science and Technology Major Project (2019SHZDZX02 to H.E.X. and 18ZR1447800 to LHZ); Shanghai Municipal Science and Technology Major Project (H.E.X.);CAS Strategic Priority Research Program (XDB37030103 to H.E.X.).

## Author Contributions

LHZ designed the expression constructs, purified the complexes, prepared the final samples for cryo-EM data collection toward the structure, participated in model building and performed structure and function data analysis, prepared figures and wrote the manuscript; LHZ prepared the cryo-EM grids, QNY and JRL performed map calculations, QNY built and refined the structure models; XHH performed structure modeling and volume calculation; QH, YMG and YL construct functional plasmids, QH performed signaling experiments under the supervision of LHZ; KW and JHS supplied material; LHZ and HEX conceived the project, wrote the manuscript.

## ADDITIONAL INFORMATION

**Competing interests:** The authors declare that they have no competing interests.

